# Olfactory receptor subgenome and expression in a highly olfactory procellariiform seabird

**DOI:** 10.1101/723924

**Authors:** Simon Yung Wa Sin, Alison Cloutier, Gabrielle Nevitt, Scott V. Edwards

**Affiliations:** Department of Organismic and Evolutionary Biology, Museum of Comparative Zoology, Harvard University, 26 Oxford Street, Cambridge, MA 02138, USA; School of Biological Sciences, The University of Hong Kong, Pok Fu Lam Road, Hong Kong SAR; Department of Neurobiology, Physiology and Behavior and the Graduate Group in Ecology, University of California, Davis, CA 95616, USA

**Keywords:** Leach’s storm petrel, *Oceanodroma leucorhoa*, sensory ecology, olfaction, olfactory receptor gene repertoire, transcriptome

## Abstract

Procellariiform seabirds are known for their well-developed olfactory capabilities, reflected by their large olfactory bulb to brain ratio and olfactory-mediated behaviors. Many species in this clade use olfactory cues for foraging and navigation, and some species can recognize individual-specific odors. Their genomes and transcriptomes may yield important clues about how the olfactory receptor (OR) subgenome was shaped by natural and sexual selection. In this study, we assembled a high-quality Leach’s storm petrel (*Oceanodroma leucorhoa*) genome to facilitate characterization of the OR repertoire. We also surveyed expressed OR genes through transcriptome analysis of the olfactory epithelium - to our knowledge, the first avian study to interrogate OR diversity in this way. We detected a large number (∼61) of intact OR genes, and identified OR genes under positive selection. In addition, we estimated that this species has the lowest proportion (∼60%) of pseudogenes compared to other waterbirds studied thus far. We show that the traditional annotation-based genome mining method underestimates OR gene number (214) as compared to copy number analysis using depth-of-coverage analysis, which estimated a total of 492 OR genes. By examining OR expression pattern in this species, we identified highly expressed OR genes, and OR genes that were differentially expressed between age groups, providing valuable insight into the development of olfactory capabilities in this and other avian species. Our genomic evidence is consistent with the Leach’s storm petrel’s well-developed olfactory sense, a key sensory foundation for its pelagic lifestyle and behavioral ecology.

## Introduction

Animals have evolved different senses to survive and flourish in changing environments. Of the several animal senses, olfaction is the physiological function detecting highly diverse and abundant chemicals originating from the surrounding environment and other organisms. Olfaction is important for animals to recognize food, mates, relatives, offspring, predators, diseases, territories, and many other important functions (Wyatt 2003). It is therefore crucial for their survival and reproduction.

In vertebrates, the ability to detect and differentiate tens of thousands of odorants is largely mediated by olfactory receptors (ORs) expressed in the olfactory epithelium of the nasal cavity (Buck and Axel 1991). Olfactory receptors are transmembrane G protein-coupled receptors (GPCRs) with seven α-helical transmembrane domains bound to a G-protein. The binding of extracellular ligands to ligand-binding sites of ORs triggers conformational changes that lead to intracellular signaling cascades, resulting in transmission to the olfactory bulb in the brain (Fredriksson, et al. 2003), which ultimately leads to olfactory perception. It has been proposed that different types of ligands are recognized by different combinations of ORs to enable an individual to perceive thousands of chemicals as distinct odors (Malnic, et al. 1999). The large number of ORs in vertebrates are classified into two groups. Class I ORs are hypothesized to bind water-borne hydrophilic ligands, and class II ORs appear to bind airborne hydrophobic ligands (Saito, et al. 2009).

Olfactory receptors are encoded by OR genes, which, at approximately 1,000 bp in size, are without introns and relatively short. OR genes are the largest multigene family in vertebrates (Nei, et al. 2008). Moreover, frequent gains and losses through duplication and pseudogenization, have resulted in dramatic differences in OR repertoire and gene number between species (Nei, et al. 2008; Nei and Rooney 2005; Niimura 2012). New OR families likely originate through gene duplication and positive selection leading to neofunctionalization and species-specific adaptions, whereas loss of function of some gene duplicates typically results in a large number of OR pseudogenes (Innan 2009; Lynch and Force 2000). The number of intact OR genes ranges from 40 in pufferfish (Niimura and Nei 2005) to ∼2000 in the African elephant (Niimura, et al. 2014). The overall size and diversity of the OR repertoire across species is believed to be influenced by ecological adaptation and reliance on olfaction (Gilad, et al. 2004; Hayden, et al. 2010). Highly olfactory mammals such as elephants have many intact OR genes compared to primate species, such as macaques, which rely more on vision than olfaction whose genomes have a smaller number of intact OR genes and a larger proportion of pseudogenes (Matsui, et al. 2010; Niimura, et al. 2014).

Among vertebrates, birds are well-known for their excellent sense of vision whereas olfaction has been largely ignored by ornithologists. However, emerging evidence shows that many birds have well-developed olfactory abilities that likely rival many mammals, including humans (Bang 1966; Bonadonna and Nevitt 2004; Corfield, et al. 2015; Nevitt, et al. 2008; Roper 1999; Zelano and Edwards 2002). The OR repertoires of birds are small relative to many other vertebrates, and gains and losses and pseudogenization seems to play an important role in their evolution (Khan, et al. 2015; Organ, et al. 2010). Ecological factors and life-history adaptations appear to have shaped the olfactory abilities and repertoire variation among birds of prey, water birds, land birds, and vocal learners (Corfield, et al. 2015; Khan, et al. 2015). Although there was an expansion in OR family 14 (the γ-c clade) in birds and the majority of avian OR genes belong to this family, some bird species and lineages exhibit alternative patterns of OR gene family expansions or reductions (Khan, et al. 2015). For example, the estimated number of OR genes is larger in the nocturnal brown kiwi (*Apteryx australis*) and flightless parrot, kakapo (*Strigops habroptilus*), than in their diurnal relatives (Steiger, et al. 2009a). In contrast, penguins, like many aquatic mammals (Hayden, et al. 2010), possess a high percentage of OR pseudogenes (Lu, et al. 2016), which appear to have been pseudogenized during the transition from a terrestrial to a marine habitat, suggesting that olfactory perception or use changed as well.

Olfactory ability is reflected by the olfactory bulb to brain ratio, which correlates positively with the estimated total number of OR genes in birds (Khan, et al. 2015; Steiger, et al. 2008). Among extant birds, the Procellariiformes, also called tube-nosed seabirds, which includes the storm-petrels, albatrosses, diving petrels, and shearwaters, have the largest olfactory bulb to brain ratio (Corfield, et al. 2015). These seabirds are known for their excellent olfactory ability. Many seabird species use olfactory cues to locate areas for foraging (Nevitt 1999a; Nevitt 2000; Nevitt 1999b; Nevitt, et al. 2004; Nevitt, et al. 1995), and several burrow-nesting species use odor to locate their burrow when returning to the colony after offshore foraging trips (Bonadonna and Bretagnolle 2002; Bonadonna, et al. 2004). Additionally, some species can recognize individual-specific odors (Bonadonna and Nevitt 2004). Olfaction therefore plays a crucial role in survival and communication in this group of seabirds. Given the importance of olfaction and the large olfactory bulb in these birds, they are good candidates for studying the evolution of avian OR genes.

Leach’s storm-petrels *Oceanodroma leucorhoa* (Vieillot, 1818), a procellariiform seabird, rely heavily on their well-developed sense of smell for foraging, homing, and mate recognition. They can smell dimethyl sulfide (DMS) and use it as a foraging cue (Nevitt and Haberman 2003). Olfaction also plays a fundamental role in social communication and individual recognition in this species (O’Dwyer, et al. 2008). Their musky smelling plumage is imbued with volatile chemicals that may give them individual olfactory signatures. They are burrow-nesting and in general adults are faithful to their burrow and mate throughout their lifetime (Morse and Buchheister 1977). In each breeding season, a breeding pair raise a single chick, which remains in the egg for 45 days and in the burrow until it fledges 60 days old to forage at sea (Warham 1990) – a remarkable life history for a bird weighing only ∼47 g. Each burrow has its own unique olfactory signature, and chicks can recognize and prefer familiar odors of their natal burrow (O’Dwyer, et al. 2008). It is suggested that a memory for familial odors may play a role later in life in the context of kin recognition and mate choice (O’Dwyer, et al. 2008). Although links to individual odor profiles have not yet been established, MHC-based mate choice by males has been recently demonstrated in this species (Hoover, et al. 2018), making it an ideal candidate for the study of OR repertoire and evolution.

Here we sequenced and assembled a high-quality genome of the Leach’s storm-petrel and characterized its OR gene family repertoire, allowing us to measure expansion and turnover in OR gene families in this procellariiform seabird and relatives. In most studies attempting to identify OR genes using genome-mining techniques such as BLAST, the sizes of OR repertoires are likely underestimated because of the collapse of similar OR sequences during assembly (Khan, et al. 2015; Sudmant, et al. 2010). We therefore also estimated the copy number (Malmstrøm, et al. 2016; Sudmant, et al. 2010) of the identified OR sequences in an effort to obtain a more accurate estimate of OR gene number. Whole-genome sequencing is the best approach to study the evolution of this large multigene family (Dehara, et al. 2012; Khan, et al. 2015; Matsui, et al. 2010; Niimura, et al. 2014; Vandewege, et al. 2016). However, at present, the northern fulmar (*Fulmarus glacialis*) is the only procellariiform species with a sequenced genome, which lacks high contiguity (contig N50 = 26k) and completeness (>10% universal single-copy orthologs missing) compared to other genomes analyzed thus far (Khan, et al. 2015). The northern fulmar genome is therefore not ideal for the identification of OR genes and estimation of OR gene copy numbers. In addition, the life-history and foraging strategies of northern fulmars are very different from Leach’s storm petrels. Northern fulmars are surface-nesting, which is a derived trait compared to most other burrow-nesting procellariiform species (van-Buskirk and Nevitt 2008). The nesting behavior has also evolved in conjunction with responsiveness to olfactory cues and foraging style (van-Buskirk and Nevitt 2008), and olfaction is likely to be the dominant sense in burrow-nesting species such as the Leach’s storm petrel.

In addition to interrogating the Leach’s storm-petrel genome, we investigated the expression of OR genes in the olfactory epithelium. Procellariiform seabirds have well-developed olfactory concha (Bang 1966) where the interaction of ORs with ligands and detection of odors takes place. However, to our knowledge, there is currently no study of OR transcriptomes in birds, including chicken. Most OR genes have been identified through comparative genomic techniques using homology searches to annotate protein coding sequences, but there is typically no experimental data to support whether identified OR genes are actually expressed in the olfactory epithelium in birds. ORs are also expressed in non-olfactory tissues (Fukuda, et al. 2004; Pluznick, et al. 2009) and in sperm (Spehr, et al. 2003). Hence it is possible that some OR genes are not expressed in olfactory epithelium and play no role in the sense of smell. In addition, the difference in expression level of different OR genes and families is unknown even for those genes that are expressed in olfactory tissues. The relationship between expression pattern and function in life-history is also important to understand olfactory-mediated behaviors. If there were sexual dimorphism or developmental differences in olfactory-mediated behaviors, OR gene expression may facilitate these differences. To study OR expression, we used transcriptome sequencing (RNA-seq) to compare OR gene expression between male and female birds, and between adults and chicks, allowing us to identify highly expressed OR genes, and OR genes differentially expressed between age classes.

## Materials and Methods

### Sample collection

We captured Leach’s storm-petrels (n = 10) at Bon Portage Island, Nova Scotia, Canada (43°26’ N, 65°45’ W), where approximately 50,000 pairs breed annually (Oxley 1999). The age class (chick or adult) and burrow number of each individual were recorded (Hoover, et al. 2018). Approximately 75 µl of blood was taken from one male via brachial venipuncture and stored in a microcentrifuge tube containing Queen’s lysis buffer (Seutin, et al. 1991) and were then stored unfrozen at 4°C until DNA extraction for whole-genome sequencing. The anterior olfactory concha and right brain were collected from three adult females, three adult males, and three chicks during August, 2015, and were stored in RNAlater at 4°C for a few days until RNA extraction. All sampling was conducted in adherence to guidelines defined by the University of California, Davis Institutional Animal Care and Use Committee Protocol #19288, and Canadian Wildlife Service (permit #SC2792).

### DNA extraction and whole-genome sequencing

We isolated genomic DNA using the DNeasy Blood and Tissue Kit (Qiagen, Hilden, Germany) and determined sex of the individual for whole-genome sequencing using published PCR primers (2550F & 2718R; Fridolfsson and Ellegren 1999). We measured DNA concentrations using a Qubit dsDNA HS Assay Kit (Invitrogen, Carlsbad, USA) and performed whole-genome libraries preparation and sequencing following Grayson et al. (2017) on an adult male. In brief, a DNA library of 220 bp insert size was prepared using the PrepX ILM 32i DNA Library Kit (Takara), and mate-pair libraries of 3 kb and 6 kb insert sizes were prepared using the Nextera Mate Pair Sample Preparation Kit (cat. No. FC-132-1001, Illumina). We then assessed library quality using the HS DNA Kit (Agilent) and quantified the libraries with qPCR prior to sequencing (KAPA library quantification kit). We sequenced the libraries on an Illumina HiSeq instrument (High Output 250 kit, PE 125 bp reads) at the Bauer Core facility at Harvard University. We assessed the quality of the sequencing data using FastQC, removed adapters using Trimmomatic (Bolger, et al. 2014), and assembled the genome using AllPaths-LG (Gnerre, et al. 2011). The completeness of the assembled genome was measured with BUSCO v2.0 (Simão, et al. 2015) and the aves_odb9 dataset to search for 4915 universal single-copy orthologs in birds.

### RNA extraction and transcriptome sequencing

RNA was extracted from each sampled tissue using RNeasy Plus Mini kit (Qiagen). The quality of the total RNA was assessed using the RNA Nano Kit (Agilent). Poly-A selection was conducted on the total RNA using the PrepX PolyA mRNA Isolation Kit (Takara). The mRNA was assessed using the RNA Pico kit (Agilent) and used to make transcriptome libraries using the PrepX RNA-Seq for Illumina Library Kit (Takara). The HS DNA Kit (Agilent) was used to assess library quality. The libraries were quantified by performing qPCR (KAPA library quantification kit) and then sequenced on a NextSeq instrument (High Output 150 kit, PE 75 bp reads). Each of a total of 29 libraries (Table S1) was sequenced to a depth of approximately 30M reads. The individuals for RNA-seq were not the same as the individual for whole-genome sequencing (Table S1).

### Genome annotation

We annotated the Leach’s storm-petrel genome using MAKER v2.31.8 (Holt and Yandell 2011). We combined *ab initio* gene prediction with protein-based evidence from 16 other vertebrates (10 birds, 3 reptiles, 2 mammals, and 1 fish species), as well as the transcriptome assembly and TopHat junctions from the Leach’s storm-petrel (Table S1). We assembled the storm-petrel transcriptome from 10 tissues of a single individual (Table S1) using TRINITY 2.1.1 (Grabherr, et al. 2011) and inferred splice junctions using TopHat 2.0.13 (Kim et al. 2013). We functionally annotated the genome to identify putative gene function and protein domains using NCBI BLAST+ and the UniProt/Swiss-Prot set of proteins. We used BLASTP on the list of proteins identified by MAKER with an evalue of 1e-6.

### Data analysis

#### OR gene identification/annotation

We identified the OR genes in the Leach’s storm-petrel genome assembly with TBLASTN searches using published intact OR amino acid sequences from Vanderwege et al. (2016), Niimura et al. (2009) and the HORDE database (The Human Olfactory Data Explorer). The queries include intact OR genes from 12 species of birds, reptiles, mammals, amphibians, and fish (Table S2). We first identified all high-scoring segment pairs (HSPs) with a minimum length of 150 bp and an e-value < 1e-10. We then used BEDTools intersect (Quinlan and Hall 2010) and custom Perl scripts to tile overlapping HSPs and remove redundant BLAST results to produce a set of candidate OR regions in the storm-petrel.

Candidate OR regions were manually reviewed to omit spurious (non-OR) hits and to determine if each region represents an intact OR gene, a pseudogene, a truncated OR sequence, or an OR gene fragment. The region spanning +/- 700 bp to each side of the predicted OR location was used in an online blastx search against the NCBI non-redundant database delimited by organism ‘Aves’. Candidate OR genes were omitted if they had top BLAST hits to non-OR sequences (e.g. other non-OR GPCRs), and coordinates for retained genes were refined based on BLAST hits to other avian ORs.

OR genes were classified as ‘intact’ if they contained start and stop codons, with no internal stops or frameshifts, and as ‘pseudogenes’ if they covered the full coding region but contained internal stops or frameshifts, or had large (> 5 amino acids) insertions or deletions within transmembrane regions. Candidate ORs that spanned incomplete coding sequences were classified as ‘truncated’ if they abutted a scaffold edge or a gap between contigs, or as an OR gene ‘fragment’ if they had an apparently naturally incomplete coding region that was not at a scaffold or contig edge. ‘Truncated’ or ‘fragmented OR genes’ could also be classified as ‘pseudogenes’ if they contained internal stops or frameshifts; OR genes could also be classified as both ‘truncated’ and ‘fragmented’ (e.g. truncated at one end and fragmented at other).

We performed a second TBLASTN search using the intact storm-petrel OR genes as queries to search back against the petrel genome assembly to identify any additional candidate regions that may have been missed in the first TBLASTN search. Candidate regions were compared to the OR genes identified in the first round of blast searching with the BEDTools subtract option, requiring 10% overlap. We then used NCBI’s conserved domain search to annotate transmembrane regions TM1-TM7.

#### Phylogenetic analysis and OR gene family assignment

We used phylogenetic analysis of OR amino acid sequences to compare intact storm-petrel OR genes to other avian and reptilian OR genes. The result was used primarily to assign Leach’s storm-petrel genes to an OR subfamily. We included intact OR sequences from the American alligator, green anole, chicken, and zebra finch from Vanderwege et al. (2016), and waterbirds, including members of Sphenisciformes, Pelecaniformes, Suliformes, Gaviiformes, Phoenicopteriformes, Podicipediformes, and Anseriformes, with assembled genomes and annotated gene models on NCBI (Jarvis, et al. 2014) (Table S3). Pseudogenes, genes encoded by multiple exons, truncated genes, and partial coding regions (< 275 AA) were omitted. We used five non-OR rhodopsin family GPCRs from chicken as outgroups (Niimura 2009). They are alpha-1A adrenergic receptor (ADRA1A), 5’ hydroxytryptamine receptor 1B (HTR1B), somatostatin receptor type 4 (SSTR4), dopamine receptor D1 (DRD1), and histamine receptor H2 (HRH2).

We aligned the sequences with the ‘einsi’ option in MAFFT v. 7.407. We manually reviewed the alignment and removed sequences with large indels (> 10 consecutive amino acids). We also removed duplicates and any sequences with > 5% uncalled residues (Xs), or > 10 Xs in total, unless they were Leach’s storm-petrel OR genes or outgroup sequences. We aligned the retained sequences again with the MAFFT einsi option as described above, following which the alignment edges were trimmed to retain only the region spanning transmembrane regions TM1-TM7 for phylogenetic analysis.

We used Prottest3 v.3.4.2 (Darriba, et al. 2011) to determine the best-fitting model of amino acid substitution, which was JTT + G + F. The best maximum-likelihood topology was inferred with RAxML v. 8.2.10 (Stamatakis 2014) from 100 searches, each starting from a different random starting tree. Five hundred bootstrap replicates were computed with RAxML, and the bootstraps were plotted on the bestML tree. The bestML + bootstraps tree was then rooted on the chicken non-OR outgroups with ETE3 (Huerta-Cepas, et al. 2016). The final tree was visualized in MEGA X (Tamura, et al. 2011). Leach’s storm-petrel genes were then assigned to an OR family based on phylogenetic relationships.

#### OR gene copy number analysis

We calculated the genomic depth-of-coverage (DoC) for each olfactory receptor gene identified in the Leach’s storm-petrel genome assembly. We then compared each DoC to the genome-wide DoC to determine if any predicted OR genes represented collapsed gene copies in the genome assembly (Malmstrøm, et al. 2016; Sudmant, et al. 2010). We could then estimate the total expected number of petrel ORs. We first repeatmasked the reference genome assembly with query species ‘vertebrata metazoa’ using RepeatMasker v. 4.0.5 (Smit, et al. 2015) with RepeatMasker Library ‘Complete Database 20160829’. The reads of the 220bp fragment libraries were trimmed with Trimmomatic v. 0.32 (Bolger, et al. 2014) and mapped to the storm-petrel genome assembly using BWA v. 0.7.15 (Li and Durbin 2010) with default parameters. SAMtools v. 1.5 (Li, et al. 2009) was used to post-process mapped reads and merge output BWA SAM files. Reads that were unmapped or below the minimum mapping quality of ‘30’ were omitted. Duplicates were marked and removed with Picard v. 2.18.9 (https://broadinstitute.github.io/picard/). Per-base depth of coverage was then output with the BEDTools v. 2.26.0 genomecov option.

To incorporate the difference in DoC due to variable GC content for the estimation of OR gene copy number, we used the repeatmasked reference genome to calculate DoC for non-repetitive regions only. We calculated DoC within bins of 1000 bp (approximately the size of an intact OR gene) with at least 98% base (non-N) occupancy. For each bin, we calculated the %GC and the average DoC. Then we calculated the mean DoC within each bin, and placed bins in categories of 5% GC (e.g. 0-5%, 5-10%, 10-15%, etc.). We took the ratio of each Leach’s storm-petrel OR gene DoC and compared it to the estimated DoC for the bins with similar GC content. This DoC analysis could not be done for other waterbirds because genome coordinates for intact, pseudo- and truncated OR genes are needed, but they are not provided in Khan et al. (2015).

#### OR gene expression analysis

We assessed the quality of the RNA-seq data using FastQC (Andrews 2010). We performed error correction using Rcorrector and removed unfixable reads using a custom python script (https://github.com/harvardinformatics/TranscriptomeAssemblyTools/blob/master/FilterUncorrectabledPEfastq.py). We next removed adapters and low quality reads (-q 5) using TrimGalore! v0.4 (Krueger 2016). We removed reads of rRNAs by mapping to the Silva rRNA database using Bowtie2 2.2.4 (Langmead and Salzberg 2012) with the --very-sensitive-local option, and retained reads that did not map to the rRNA database.

We used RSEM (v1.2.29) (Li and Dewey 2011) to quantify levels of gene expression. We first built an RSEM index for the annotated Leach’s storm-petrel genome, then used RSEM to implement Bowtie2 (v2.2.6) for the mapping of RNA-seq reads to the genome, using default parameters for mapping and expression quantification. Expected read counts per million at the gene level from RSEM were used to represent the normalized expression. We used the normalized counts rounded from RSEM outputs as inputs for differential expression analysis. We then used limma voom (Law, et al. 2014) to identify differentially expressed genes between adults and chicks, and between male and female adults, using a 5% FDR cutoff.

#### Gene ontology (GO) analysis

We used GOrilla to perform GO analysis (Eden, et al. 2009), using the single ranked list of genes mode. Reported enrichment *p* values were FDR-adjusted using the Benjamini– Hochberg method (Benjamini and Hochberg 1995).

#### Analysis of positive selection on OR family 14

We detected sites that were under selection by investigating the ratio of the rate of synonymous substitutions to the rate of non-synonymous substitutions (*ω* = d*N*/d*S*), which may indicate positive selection (*ω* > 1), neutral (*ω* = 1), or negative selection (*ω* < 1). We used the HyPhy package (Pond and Muse 2005) implemented in the Datamonkey webserver (datamonkey.org) to infer potential recombination breakpoints and estimate *ω*. Since recombination and gene conversion can mislead estimation of selection, we used Genetic Algorithm for Recombination Detection (GARD) (Pond, et al. 2006) to generate multiple phylogenies based on putative non-recombinant fragments. We then used Single-Likelihood Ancestor Counting (SLAC), Fixed Effects Likelihood (FEL), Mixed Effects Model of Evolution (MEME), and Fast Unconstrained Bayesian AppRoximation (FUBAR) methods implemented in HyPhy, plus an integrated approach that incorporates all sites detected by each method, to infer signals of positive selection. Here, sites detected by two or more methods are considered under selection. All methods were used with default settings. We used WebLogo (weblogo.threeplusone.com) to visualize the amino acid sequence variation of the transmembrane (TM), intracellular (IC) and extracellular (EC) domains.

## Results

### Assembly of Leach’s storm-petrel genome

We generated 439,914,448 reads from the 220 bp library, 313,504,024 reads from the 3 kb library, and 269,594,574 reads from the 6 kb library. The genome size estimated by AllPaths-LG from k-mers is 1.24 Gb (Table 1). The contig N50 is 165.4 kb and the scaffold N50 is 8.7 Mb (Table 1). BUSCO (Simão, et al. 2015) shows a high completeness of the genome, with 98.0% of single-copy orthologs for birds identified and 94.7% represented by complete coding sequences in the genome (Table 1). The MAKER run identified a total of 15510 gene models. The genome-wide GC content is 42.1%.

**Table 1.** Assembly statistics for Leach’s storm petrel genome

|  | Leach's storm petrel genome |
| --- | --- |
| Estimated genome size | 1.24 Gb |
| %GC content | 42.1 |
| Total depth of coverage | 80x |
| Total contig length (bp) | 1,181,786,487 |
| Total scaffold length (bp, gapped) | 1,195,165,757 |
| Number of contigs | 17396 |
| Contig N50 (bp) | 165.4 kb |
| Number of scaffolds | 1697 |
| Scaffold N50 (with gaps) | 8.7 Mb |
| Total BUSCOs | 4817/4915 (98.0%) |
| Complete BUSCOs | 4654/4915 (94.7%) |

### OR genes in Leach’s storm petrel

We identified 221 candidate OR regions from the initial round of TBLASTN (Table 2). Eight of these regions were not ORs. The second TBLASTN search using all identified intact OR genes as queries identified one additional pseudogene fragment region not found in the initial round of search, yielding 214 OR regions in total. Of these 214 OR regions, 61 (28.5%) were intact OR genes, and the remainder included 106 pseudogenes (49.5%), 20 truncated genes (9.3%), and/or 27 gene fragments (12.6%) (Table 2; Fig. 1).

**Figure 1.**
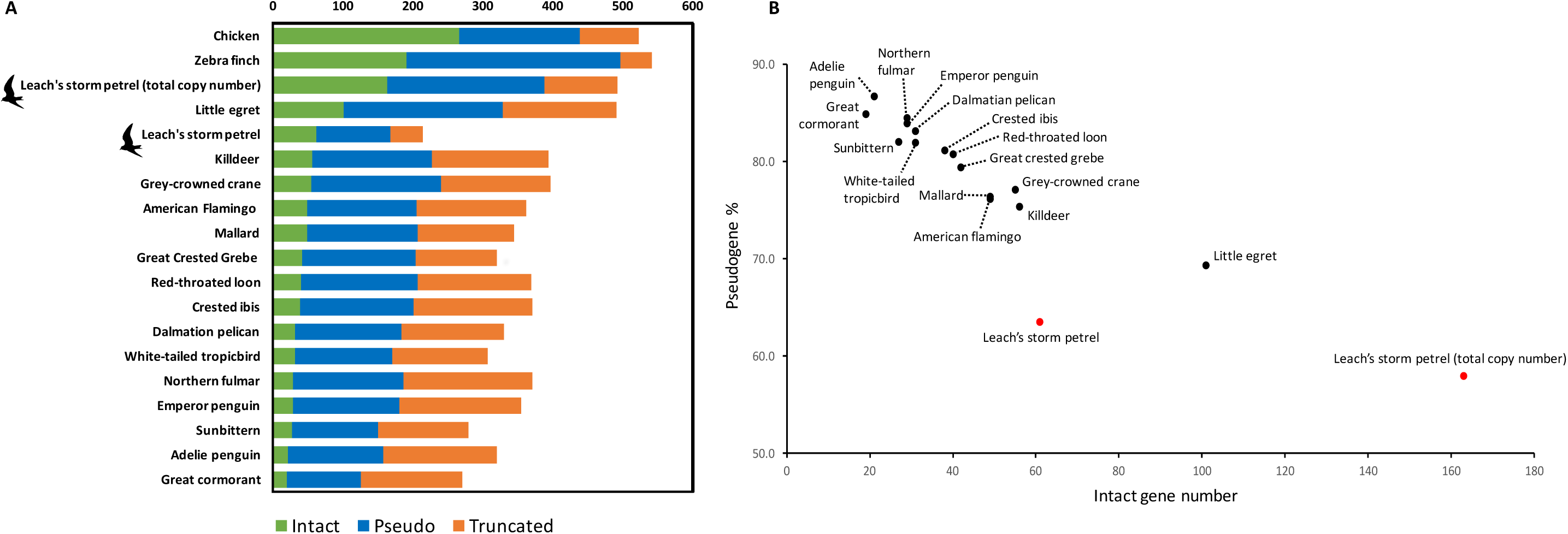
A) The number of truncated, pseudo-, and intact OR genes in waterbirds, chicken, and zebra finch. B) The number of intact genes plotted against the percentage of pseudogenes within the same genome in waterbirds. Both the OR gene number estimations based on genome annotation and copy number calculation in the Leach’s storm petrel are shown. The numbers for all species except the Leach’s storm-petrel are from Khan *et al*. (2015).

**Table 2.** The number of intact, pseudo-, truncated, and fragment OR genes and their average coverage in the Leach’s storm petrel genome.

|  | Number of<br>genes | Total copy<br>number <sup>A</sup> | Average<br>coverage |
| --- | --- | --- | --- |
| **Intact** | **61** | **163** | **2.7** |
| **Truncated** | **20** | **51** | **2.6** |
| **Total pseudogene** | **106** | **224** | **2.1** |
| • Pseudogene | • 45 | • 81 |  |
| • pseudogene/fragment | • 49 | • 103 |  |
| • pseudogene/fragment/truncated | • 2 | • 4 |  |
| • pseudogene/truncated | • 10 | • 36 |  |
| **Total fragment** | **27** | **54** | **2** |
| • fragment | • 24 | • 48 |  |
| • fragment/truncated | • 3 | • 6 |  |
| **Total (I+T+P+F)** | **214** | **492** | **2.3** |

<sup>A</sup> Refer to the Discussion for the limitation of copy number estimation.

To estimate the total number of OR genes, we incorporated the number of collapsed gene copies for the 214 identified OR genes. By calculating the ratio of each OR gene DoC to the estimated DoC for bins of similar GC content across the storm-petrel genome (Fig. S1), we estimated there are as many as 492 predicted OR genes in the Leach’s storm-petrel genome (Table 2). As expected, genes in high GC bins (> 50% GC) had lower coverage than genes in low GC bins (< 45%; Botero-Castro, et al. 2017). The average estimated copy number for intact OR genes was 2.7 and the total number of intact OR genes was 163 (33.1%) (Table 2). The copy number of intact OR genes ranged from 1 to 45 (mean = 2.7, SD = 5.8) (Table S4). Of the 24 intact OR genes with multiple copies, 13 belonged to OR family 14 (γ-c clade; Khan, et al. 2015), which included the intact gene with the highest copy number ratio of 45. The total number of estimated pseudogenes, truncated genes, and gene fragments was 224 (45.5%), 51 (10.4%), and 54 (11%), respectively (Table 2; Fig. 1).

### OR gene family phylogeny

We performed phylogenetic analyses using all intact OR genes from the Leach’s storm-petrel genomes and 13 waterbirds, plus ORs from American alligator, green anole, chicken, and zebra finch. We found that sequences largely cluster by OR gene family, although typically with low bootstrap support. Nevertheless, we were able to confidently assign 60 of 61 intact storm-petrel ORs to their OR gene family. The resulting phylogeny implied 10 OR gene families in the Leach’s storm-petrel genome (Fig. 2), corresponding to numbers 2, 4, 5, 6, 8, 10, 13, 14, 51, and 52 in chicken.

**Figure 2.**
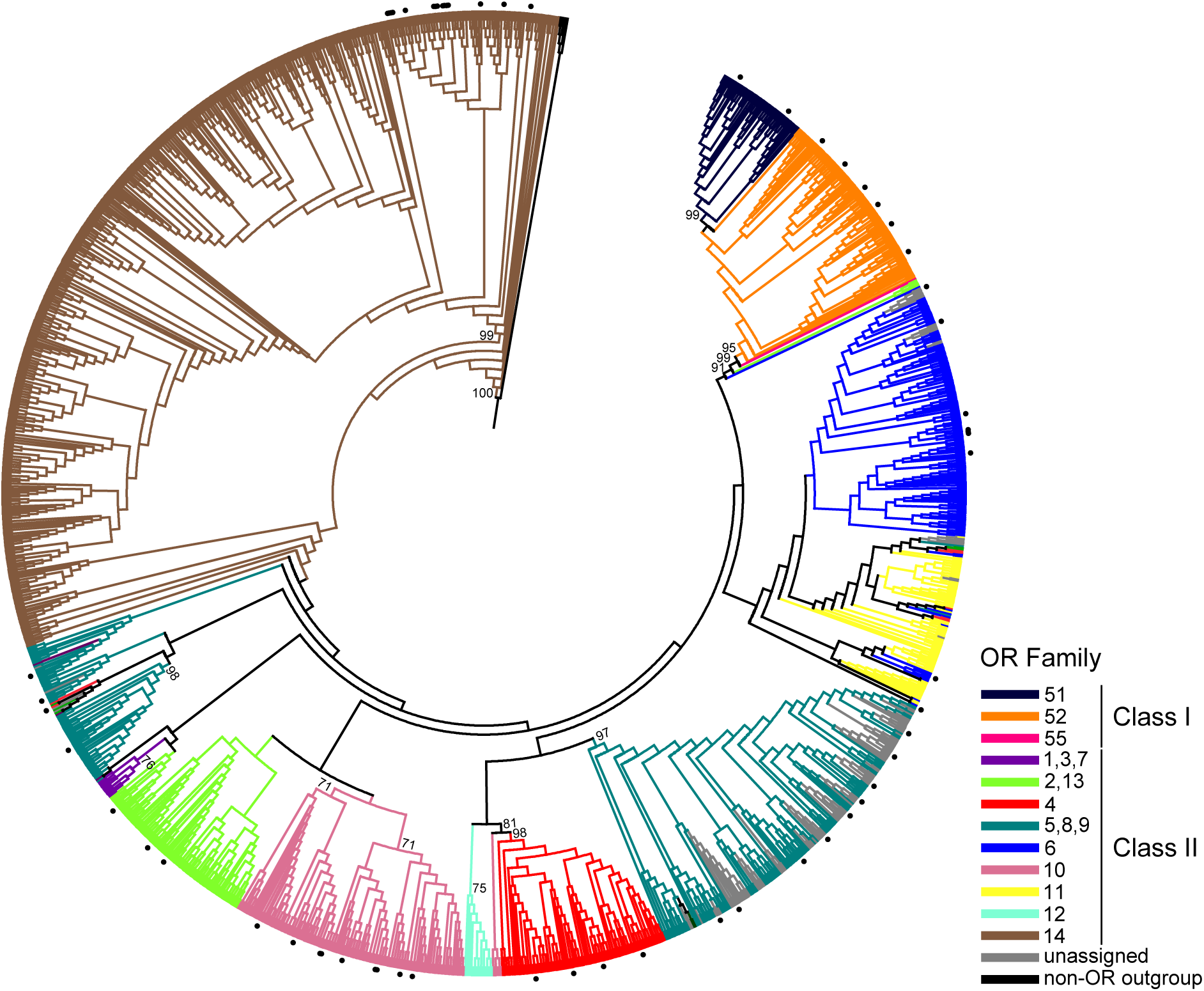
Maximum-likelihood topology of relationships among intact ORs. Branches for individual OR sequences are coloured according to OR family, and branch lengths are not drawn to scale. Circle symbols indicate intact OR genes identified in Leach’s storm petrel. Percentage support values from 500 bootstrap replicates are indicated for major clades with > 70% support.

### Differential OR gene expression

We compared the patterns of OR gene expression in the olfactory concha, where OR genes are expected to be predominantly expressed, and in the brain, where we expect little OR gene expression (Fig. 3). Two OR genes were highly expressed in the olfactory epithelium: OR gene OR6-6 (OR family 6) and OR5-11 (OR family 5). Both OR genes had a copy number ratio of two. We found no differentially expressed OR genes in the olfactory epithelium between male and female adults (Fig. S2), but identified four OR genes differentially expressed between age classes: OR14-14, OR14-12, OR10-2, and OR14-9 (Fig. 3). The most differentially expressed OR gene, OR14-14, is also the OR gene with the highest copy number ratio at 45 (Table S4). OR14-12 and OR14-9 also had a relatively high copy number ratio at 5 and 9, respectively (Table S4). The two highly expressed ORs and four differentially expressed ORs are all class II ORs. In contrast to the expression in the olfactory epithelium, most OR genes were not expressed or exhibited minimal (∼0) expression in the brain (Fig. 3C), and there were no differentially expressed OR genes in the brain sample. Gene ontology (GO) analyses of 6101 genes significantly differentially expressed (FDR < 0.01) in the olfactory epithelium between age classes revealed categories related to tissue growth and development, such as ossification and collagen fibril organization, as the most significantly enriched (Table S5). There were only 28 genes differentially expressed between adult males and females in the olfactory epithelium, with no GO categories enriched.

**Figure 3.**
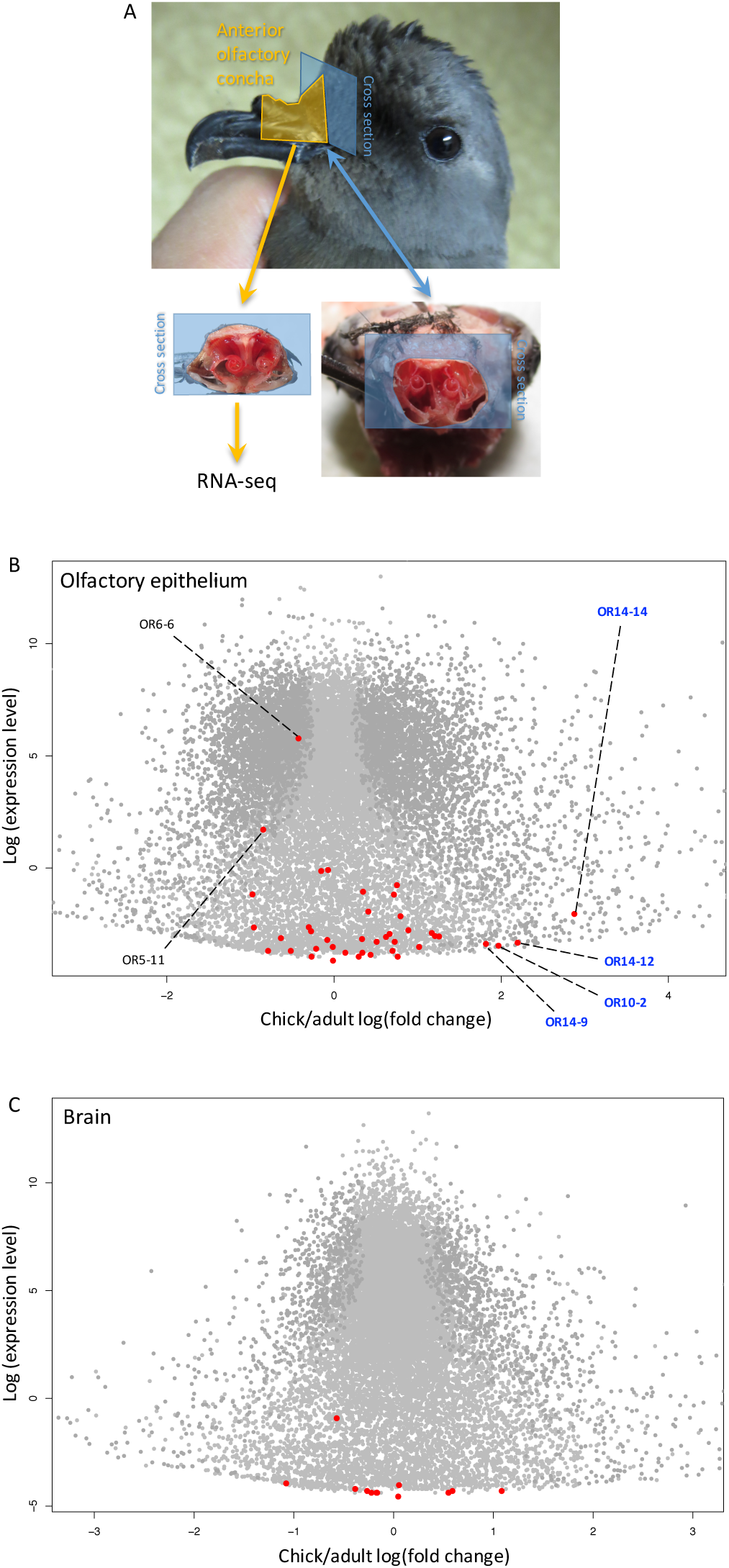
OR genes expression in the olfactory epithelium of anterior olfactory concha of the Leach’s storm petrel. A) Anterior olfactory concha of the Leach’s storm petrel for RNA-seq. B) Differentiation expression of the genes in chick versus adult olfactory epithelium. Differentially expressed genes are in dark grey. OR genes are highlighted in red. Four OR genes with higher expression in chicks are labelled with their names in blue. Two most highly expressed OR genes are also labelled. C) Differentiation expression of the genes in chick versus adult brain. No OR genes were differentially expressed in the brain.

### OR genes under positive selection

We found evidence of two recombination breakpoints at nucleotide position 321 and 450 of the alignment, located in the TM3 and TM4 domains, respectively (Fig. 4). Based on the inferred breakpoints, we used three data partitions to identify sites under selection in the intact genes of OR family 14. We identified signals of positive selection in OR family 14 using multiple approaches. Although the overall *ω* was 0.449 (SLAC), 0.436 (FEL), and 0.449 (MEME), which suggest no evidence of positive selection across the genes as a whole, we detected signals of positive selection in individual codons. We identified codon positions 4 and 107 (in TM3 domain) to be under positive selection using all methods (Table 3; Fig. 4). Codon positions 156 (in TM4), 200 (in TM5), and 250 (in TM6) were also under positive selection, identified by at least two methods (Table 3; Fig. 4).

**Table 3.**
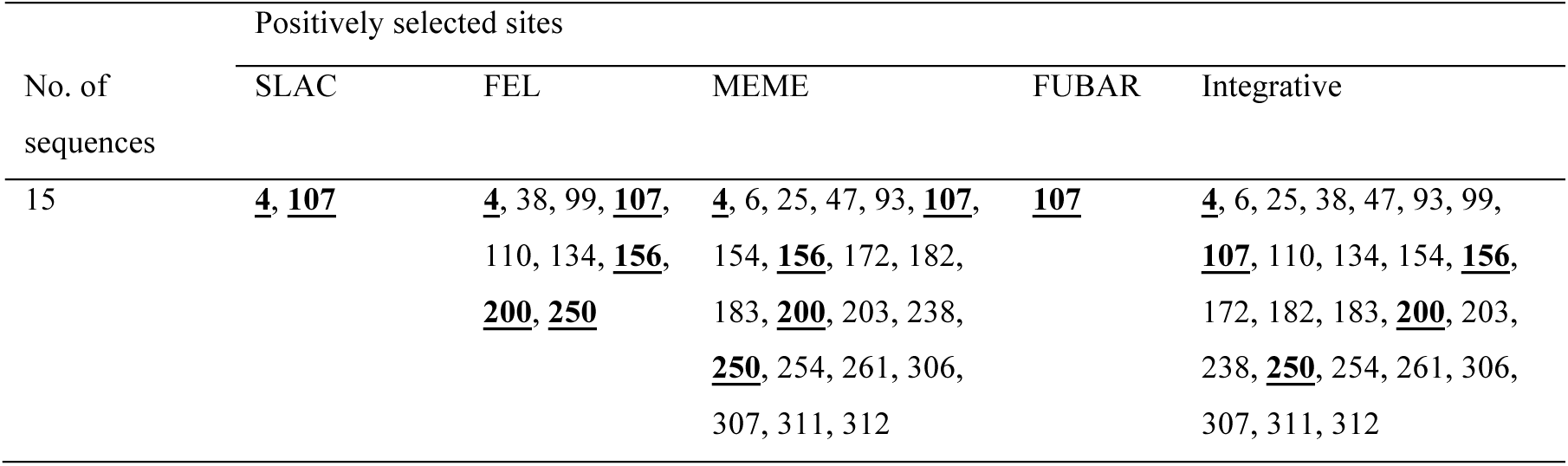
Positively selected sites detected by five approaches, along with integrated analysis, in genes of OR family 14 in the Leach’s storm petrel. The sites detected by more than two methods are in bold and underlined.

**Figure 4.**
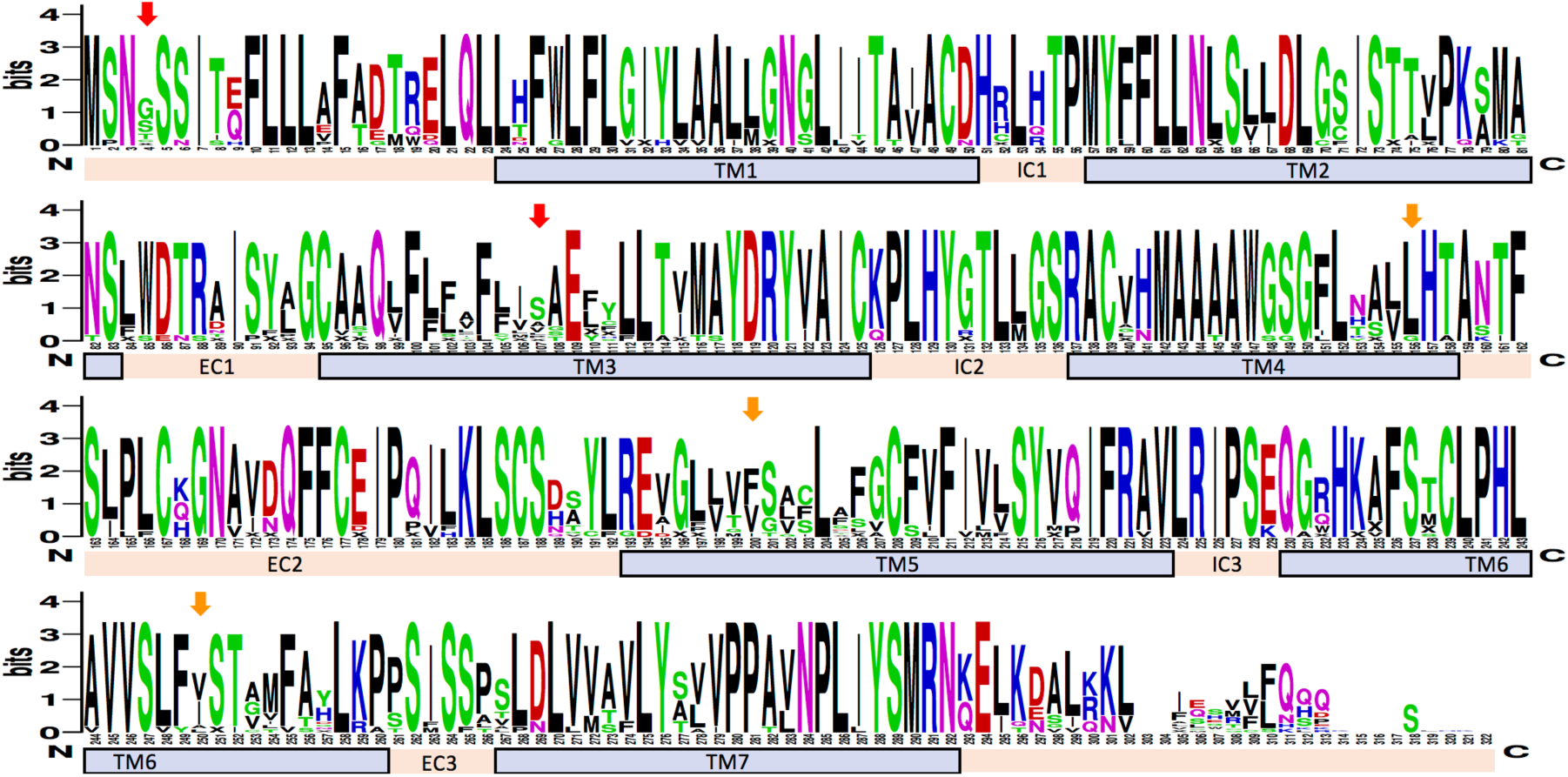
Amino acid sequence variation of the intact family 14 OR genes in the Leach’s storm petrel. Red and orange arrows indicate significant positively selected sites identified by all and at least two methods, respectively. Locations of the transmembrane domains (TM1-7), intra-cellular domains (IC1-3), and extra-cellular domains (EC1-3) are shown. The overall height of the stack of symbols indicates the sequence conservation at that codon position. The height of amino acid symbols with the stack indicates the relative frequency of each amino acid at that codon position. Numbers below the stacks indicate codon position.

## Discussion

Our high-quality genome of a Leach’s storm-petrel has higher contiguity than many bird genomes produced with short-read technology and allowed us to identify 61 intact OR genes and to estimate the proportion of intact and pseudogenized ORs. Because highly similar sequences from short-read libraries often lead to misassembled genes during whole-genome assembly (Alkan, et al. 2011), we examined the copy number ratio of OR sequences using depth-of-coverage and estimated a more than two-fold increase in OR gene number as compared to the annotation-only method. The OR gene number estimate incorporating the copy number ratio should be closer to the actual number of OR genes in this species (Malmstrøm, et al. 2016; Sudmant, et al. 2010). The actual number of OR genes is probably underestimated in most studies using genome blast-based mining and annotation only method to identify highly similar duplicated genes, a situation similar to the case of highly duplicated MHC genes (Malmstrøm, et al. 2016). Mapping of sequencing reads to estimate the copy number ratio is one way to better estimate the actual gene copy number (Malmstrøm, et al. 2016). A limitation of this approach, however, is that the sequencing reads are usually shorter than the assembled OR sequences in the reference genome, and the highly similar nature of OR sequence also makes mapping assignment difficult or impossible, therefore the mapping depth-of-coverage for each OR sequence may deviate from the actual copy number ratio. Nonetheless, the total OR copy number estimate should be more accurate using this approach than genome mining alone. To provide a more accurate copy number estimation in the future, the emerging strategies using long-read sequencing technology that generates tens of kilobases read length can aid the study of multigene families such as OR genes (Miller, et al. 2017).

When compared to other waterbirds (Khan, et al. 2015), the number of intact genes in Leach’s storm-petrels is the highest if we consider the estimated copy number (Fig. 1). It is also among the highest in intact OR number even when estimates of copy number are not considered, and is less than only one waterbird, the little egret (Fig. 1). The proportion of pseudogenes (pesudogene/(pseudogene+intact gene)) is the lowest among waterbirds, at approximately 60% in the Leach’s storm-petrel compared to 69%-87% in other waterbirds (Fig. 1). Despite being the sister group to the Procellariiformes, the penguins (Sphenisciformes), represented here with Adelie and emperor penguins, are among the species with the lowest number of intact genes and the highest proportion of pseudogenes. This pattern may be due to their obligate mode of foraging underwater via diving behavior (Lu, et al. 2016). Another procellariiform seabird, the northern fulmar, has a similarly low number of intact genes and high proportion of pseudogenes as in penguins. One possibility is that the sequencing depth and genome assembly quality of the northern fulmar is much lower than that of the Leach’s storm-petrel sequenced here, because the number of OR genes identified in the chicken and zebra finch, which have high-quality assembled genomes, was larger. However, the quality of the fulmar genome is comparable to many other waterbird genomes, and Khan et al. (2015) showed that there was no correlation between the number of OR genes identified and genome-wide sequencing depth. The high OR gene number in chicken (266 intact genes; 39.4% pseudogene) and zebra finch (190 intact genes; 61.7% pseudogene) may be due to species- or lineage-specific expansion in these groups (Fig. S3) (Khan, et al. 2015). Perhaps the different OR repertoires of the Leach’s storm petrel and other waterbirds is a real biological signal that arose during the diversification of OR genes in different bird lineages.

The larger number of intact OR genes and smaller percentage of pseudogenized ORs in Leach’s storm-petrels than most waterbirds suggests enhanced olfactory capabilities, consistent with the large olfactory bulb ratio in Procellariiformes (Corfield, et al. 2015; Khan, et al. 2015; Steiger, et al. 2008), and is supported by behavioral tests revealing a well-developed sense of smell in this species (Nevitt and Haberman 2003; O’Dwyer, et al. 2008). Gene gains and losses through gene duplication and pseudogenization are the main processes in OR evolution among birds and other vertebrates (Khan, et al. 2015; Lu, et al. 2016; Niimura, et al. 2014; Steiger, et al. 2009b). The use of olfaction for behaviors such as foraging, homing, and mate recognition in the Leach’s storm-petrel could be the selective force driving the evolution of OR gene number in this species. Being exclusively pelagic, procellariforms are adapted to forage efficiently in order to survive in a vast area of open ocean where food sources can be patchy, unpredictable and transient. Explanation of OR gene number involving foraging habitat is not universal when we consider the low number of intact ORs in the northern fulmar, the only other Procellariiform with its genome sequenced and OR genes studied. There is a diversity of behaviors and ecologies among Procellariiform species. For example, Leach’s storm petrels incubate their eggs and feed their chicks inside an underground burrow, whereas the northern fulmar nest on the ground (van-Buskirk and Nevitt 2008). Leach’s storm petrel chicks spend almost the entire nestling period underground before they fledge, and parents enter and exit the breeding colony to feed their chicks nocturnally, when predation risk is lower. This difference in rearing environment could lead to differences in sensory functions (van-Buskirk and Nevitt 2008). A strong reliance on olfaction and good sense of smell may develop in Leach’s storm-petrels being raised in darkness, whereas Procellariiform species exposed to more light may depend less on olfaction for homing and individual recognition (Mitkus, et al. 2016; Mitkus, et al. 2018). The Leach’s storm-petrel indeed has six times lower visual spatial resolution than the northern fulmar (Mitkus, et al. 2016), which rely more on using vision than olfaction for foraging (van-Buskirk and Nevitt 2008). By investigating the OR subgenome in this study, our genomic and transcriptomic evidence confirms that the Leach’s storm-petrel has superior olfactory capabilities among waterbirds and birds in general. Future studies should focus on the relationship between OR repertoire and species-specific behavioral ecology in a wider and more densely sampled phylogenetic context to understand how natural and sexual selection shapes avian OR evolution.

Although the phylogenetic analysis did not reveal obvious species-specific expansion of a particular OR gene family in this species (but we cannot rule this out because we do not know if the highly duplicated gene copies would have orthologs in other species), several OR genes and domains experienced positive selection. We identified five amino acid sites under positive selection on OR family 14, the family that underwent rapid expansion in birds and showed signals of positive selection in eight other bird species (Khan, et al. 2015). Four of the five positively selected sites were located in transmembrane domains 3, 4, 5, and 6. These regions were also found to be highly variable in other species, and were suggested to participate in ligand binding (Niimura 2012; Quignon, et al. 2005). Specific genes belonging to OR family 14 had a high copy number when we examined the depth of coverage. This family belongs to class II ORs that bind airborne hydrophobic ligands and probably play a crucial role in the olfactory sense of this species, given the high number of copies in the genome.

OR genes experiencing substantial duplications, in particular OR14-14, suggest their high relevance to the ecology of Leach’s storm-petrel. Identification of specific ligands for these ORs will help clarify the driving force for increasing gene copy number. For example, they may be important for foraging if OR14-14 or other OR 14-family genes bind dimethyl sulfide (DMS) or other ligands used in foraging (Nevitt, et al. 1995), or for communication and recognition if they bind odorants produced by other individuals. It is well known that individual olfactory sensory neurons (OSN) express a single OR allele out of hundreds of loci and alleles in the genome (Khamlichi and Feil 2018; Monahan and Lomvardas 2015). This monoallelic expression of OR genes determines the olfactory sensitivity of the neuron, determining the ligands that will stimulate it. The single expressed OR also instructs axonal connections of the OSN to a specific glomerulus in the olfactory bulb. Expression of more than one OR allele may lead to disruption of olfactory network wiring and misinterpretation of the sense of smell (Magklara and Lomvardas 2013). The expression mechanism of a single OR per neuron is stochastic, initiated by random chromatin-mediated activation of a single OR expression and a feedback loop that stabilizes the initial OR and prevents additional OR allele expression (Chess 2012; Eckersley-Maslin and Spector 2014; Magklara and Lomvardas 2013). Under this random monoallelic expression, an OR gene with more copies in the genome should have a larger representation in the OSN population than OR genes with a low copy number. Decoding and deorphanizing those highly duplicated ORs is a fascinating area for future research linking the olfactory environment, behavior and OR evolution.

To confirm that the identified intact OR genes are actually expressed in the olfactory epithelium we studied the transcriptome of the anterior olfactory concha. The intact OR genes identified transcriptomically were expressed in the olfactory epithelium, and different ORs were expressed at different levels. OR expression was almost absent in the brain sample, which likely included several subportions of the storm-petrel brain, including the olfactory bulb. The pattern of OR expression supports the role of identified OR genes in the detection of smell. To our knowledge, ours is the first study to investigate OR expression in the olfactory epithelium of birds using a transcriptomic approach. In other studies, once OR genes are identified by genome mining methods, there is often little confirmation to support the expression of OR genes in the olfactory epithelium. Interpreting OR gene evolution and understanding their relevance to sensory behavior may be hampered by the assumption that all annotated OR genes play a role in the sense of smell. By determining the expression of OR genes in different body tissues, we will be able to refine the functional interpretation of different OR genes, which may have roles outside of smell (Fukuda, et al. 2004; Pluznick, et al. 2009; Spehr, et al. 2003). The differences in OR expression level among OR genes could be due to spatial patterning of OSN types in the olfactory epithelium (Coleman, et al. 2019). Now that we have identified the OR genes and transcripts in this study, future investigations can focus on the spatial and temporal patterns of OR gene expression, which is a research area currently lacking in birds, and has only been studied in a few non-avian model species such as mice (Coleman, et al. 2019; Hanchate, et al. 2015).

We found four OR genes that were differentially expressed in the olfactory epithelium between adults and chicks, belonging to families 14 and 10, both of which are class II ORs. All four genes were more highly expressed in chicks. Leach’s storm-petrels can readily perform odor discrimination tasks as chicks soon after hatching (O’Dwyer, et al. 2008). A recent study by Mitkus et al. (2018) has shown that Leach’s storm-petrel chicks are blind for the first 2 to 3 weeks post hatching suggesting a heightened reliance on olfaction. In our study, some of the most over-expressed genes we identified in chick compared to adult olfactory conchae are those that involved in ossification and soft tissue development (Table S5), such as the genes *SPARC*, *PHOSPHO1*, *Smpd3*, *COL1A1*, *COL1A2*, and *COL11A1*. The olfactory epithelium, as well as the sense of smell, of chicks sampled here were probably developing rapidly when sampled, perhaps resulting in higher expression levels of some OR genes in chicks than in adults. Alternatively, the lifespan of OSNs is affected by how frequently the ORs are used (Santoro and Dulac 2012). There is a mechanism to reduce the lifespan of OSNs that express infrequently used ORs (Santoro and Dulac 2012). This process can modulate the OSN population dynamics to adapt the olfactory system to a particular environment by changing the relative number of different types of OSNs, and the relative abundance of different OSNs changes with age and experience (Santoro and Dulac 2012; van-der-Linden, et al. 2018). Adult storm petrels that are foraging, navigating, homing, and recognizing mates, likely express a different repertoire of ORs than developing chicks, which spend their entire early life inside their home burrows (but they are also interacting with the adults, feeding, walking around inside the burrow and they are very capable of discriminating different types of odors in choice tests). The difference in OR expression between chicks and adults might be caused by the difference in the usage frequency of different type of ORs, leading to variation in the lifespan and abundance of each type of OSN.

It has been proposed that MHC genes can affect body odor by changing the peptide community in the body (Brennan and Zufall 2006; Restrepo, et al. 2006). Alternatively, or in addition, individuals with different MHC genotypes may harbor different microbiome, which in turn produce different secondary metabolites and odor (Pearce, et al. 2017; Zomer, et al. 2009). Highly diverse bacterial communities are often found in animal scent glands (Sin, et al. 2012; Theis, et al. 2013), and the uropygial gland of birds is one potential place that the secretion odor is affected by the microbiome it harbors (Rodríguez-Ruano, et al. 2015; Whittaker, et al. 2016). In Leach’s storm-petrels, males appear to select their mates based on the MHC genotypes but females do not (Hoover, et al. 2018). Some insects using odors to select mates exhibit sexual dimorphism in the olfactory system (Brand, et al. 2018). The sex difference in MHC-based mate choice behavior in this species might be mediated through differentiated olfactory response to candidate mates with different body odors, which in turn could be due to intersexual differences in olfactory capabilities. An effect of MHC genes on body odor is yet to be shown in this species. Our study of gene expression in the olfactory epithelium revealed no intersexual differences in OR expression in adults. Thus, our study does not support the idea that intersexual differences in MHC-based mate choice behavior were due to different OR gene usages. However, this does not rule out that sexual dimorphism occurs in the olfactory center of the brain. Future studies of the relationship between MHC genotypes and body odor, and behavioral responses of birds to odors from birds of different MHC genotypes, could help clarify whether mate choice in this species is mediated by olfaction.

## Author contributions

S.Y.W.S, G.N. and S.V.E. designed research; S.Y.W.S. performed research; S.Y.W.S. and A. C. analyzed data; S.Y.W.S. wrote the paper and all authors contributed to revised versions.

## Acknowledgements

This research was supported by NSF (award numbers: NSF Grant IOS-1258784, NSF IOS 0922640/IBN 0212467 and NSF Grant IOS 1258828). We thank Lee Adams and David Shutler for logistical support, Marcel Losekoot for data management, Brian Hoover and Logan Lewis-Mummert for field assistance at UC Davis, Prof. Shelley Adamo and Laura Hall at Dalhousie University and the Bauer Core Facility at Harvard University (especially Jennifer Couget, Christian Daly and Claire Reardon) for laboratory assistance. We thank Tim Sackton for his help with genome assembly. The computations in this paper were performed on the Odyssey cluster at Harvard University and supported by Harvard University Research Computing.

## Data Accessibility

The draft genome and transcriptomic data are available via Dryad (DOI will be provided later).

The authors declare no conflict of interest.

## References

Alkan C, Sajjadian S, Eichler EE 2011. Limitations of next-generation genome sequence assembly. Nature Methods 8: 61.

Andrews S 2010. FastQC: a quality control tool for high throughput sequence data. Available online at: http://www.bioinformatics.babraham.ac.uk/projects/fastqc.

Bang B 1966. The olfactory apparatus of tubenosed birds (Procellariiformes). Cells Tissues Organs 65: 391–415.

Benjamini Y, Hochberg Y 1995. Controlling the false discovery rate: a practical and powerful approach to multiple testing. Journal of the Royal statistical society: series B (Methodological) 57: 289–300.

Bolger A, Lohse M, Usadel B 2014. Trimmomatic: a flexible trimmer for Illumina sequence data. Bioinformatics 30: 2114–2120.

Bonadonna F, Bretagnolle V 2002. Smelling home: a good solution for burrow-finding in nocturnal petrels? Journal of Experimental Biology 205: 2519–2523.

Bonadonna F, Nevitt GA 2004. Partner-specific odor recognition in an Antarctic seabird. Science 306: 835.

Bonadonna F, Villafane M, Bajzak C, Jouventin P 2004. Recognition of burrow’s olfactory signature in blue petrels, *Halobaena caerulea*: an efficient discrimination mechanism in the dark. Animal Behaviour 67: 893–898.

Botero-Castro F, Figuet E, Tilak M, Nabholz B, Galtier N 2017. Avian genomes revisited: hidden genes uncovered and the rates versus traits paradox in birds. Molecular Biology and Evolution 34: 3123–3131.

Brand P, Larcher V, Couto A, Sandoz J, Ramírez S 2018. Sexual dimorphism in visual and olfactory brain centers in the perfume-collecting orchid bee *Euglossa dilemma* (Hymenoptera, Apidae). Journal of Comparative Neurology 526: 2068–2077.

Brennan P, Zufall F 2006. Pheromonal communication in vertebrates. Nature 444: 308.

Buck L, Axel R 1991. A novel multigene family may encode odorant receptors: a molecular basis for odor recognition. Cell 65: 175–187.

Chess A 2012. Mechanisms and consequences of widespread random monoallelic expression. Nature Reviews Genetics 13: 421.

Coleman J, et al. 2019. Spatial determination of neuronal diversification in the olfactory epithelium. Journal of Neuroscience 39: 814–832.

Corfield J, et al. 2015. Diversity in olfactory bulb size in birds reflects allometry, ecology, and phylogeny. Frontiers in Neuroanatomy 9: 102.

Darriba D, Taboada G, Doallo R, Posada D 2011. ProtTest 3: fast selection of best-fit models of protein evolution. Bioinformatics 27: 1164–1165.

Dehara Y, et al. 2012. Characterization of squamate olfactory receptor genes and their transcripts by the high-throughput sequencing approach. Genome Biology and Evolution 4: 602–616.

Eckersley-Maslin M, Spector D 2014. Random monoallelic expression: regulating gene expression one allele at a time. Trends in Genetics 30: 237–244.

Eden E, Navon R, Steinfeld I, Lipson D, Yakhini Z 2009. GOrilla: a tool for discovery and visualization of enriched GO terms in ranked gene lists. BMC Bioinformatics 10: 48.

Fredriksson R, Lagerström M, Lundin L, Schiöth H 2003. The G-protein-coupled receptors in the human genome form five main families. Phylogenetic analysis, paralogon groups, and fingerprints. Molecular Pharmacology 63: 1256–1272.

Fridolfsson AK, Ellegren H 1999. A simple and universal method for molecular sexing of non-ratite birds. Journal of Avian Biology 30: 116–121.

Fukuda N, Yomogida K, Okabe M, Touhara K 2004. Functional characterization of a mouse testicular olfactory receptor and its role in chemosensing and in regulation of sperm motility. Journal of Cell Science 117: 5835–5845.

Gilad Y, Wiebe V, Przeworski M, Lancet D, Pääbo S 2004. Loss of olfactory receptor genes coincides with the acquisition of full trichromatic vision in primates. PLoS Biology 2: 120–125.

Gnerre S, et al. 2011. High-quality draft assemblies of mammalian genomes from massively parallel sequence data. Proceedings of the National Academy of Sciences 108: 1513–1518.

Grabherr MG, et al. 2011. Full-length transcriptome assembly from RNA-Seq data without a reference genome. Nature Biotechnology 29: 644–652.

Grayson P, Sin S, Sackton T, Edwards S. 2017. Comparative genomics as a foundation for evo-devo studies in birds: Humana Press, New York, NY.

Hanchate N, et al. 2015. Single-cell transcriptomics reveals receptor transformations during olfactory neurogenesis. Science 350: 1251–1255.

Hayden S, et al. 2010. Ecological adaptation determines functional mammalian olfactory subgenomes. Genome Research 20: 1–9.

Holt C, Yandell M 2011. MAKER2: an annotation pipeline and genome-database management tool for second-generation genome projects. BMC Bioinformatics 12: 491.

Hoover B, et al. 2018. Ecology can inform genetics: Disassortative mating contributes to MHC polymorphism in Leach’s storm-petrels (*Oceanodroma leucorhoa*). Molecular Ecology 27: 3371–3385.

Huerta-Cepas J, Serra F, Bork P 2016. ETE 3: reconstruction, analysis, and visualization of phylogenomic data. Molecular Biology and Evolution 33: 1635–1638.

Innan H 2009. Population genetic models of duplicated genes. Genetica 137: 19.

Jarvis E, et al. 2014. Whole-genome analyses resolve early branches in the tree of life of modern birds. Science 346: 1320–1331.

Khamlichi A, Feil R 2018. Parallels between mammalian mechanisms of monoallelic gene expression. Trends in Genetics 34: 954–971.

Khan I, et al. 2015. Olfactory receptor subgenomes linked with broad ecological adaptations in Sauropsida. Molecular Biology and Evolution 32: 2832–2843.

Krueger F 2016. Trim Galore. Babraham Bioinformatics.

Langmead B, Salzberg SL 2012. Fast gapped-read alignment with Bowtie 2. Nature Methods 9: 357–359.

Law C, Chen Y, Shi W, Smyth G 2014. voom: Precision weights unlock linear model analysis tools for RNA-seq read counts. Genome Biology 15: R29.

Li B, Dewey C 2011. RSEM: accurate transcript quantification from RNA-Seq data with or without a reference genome. BMC Bioinformatics 12: 323.

Li H, Durbin R 2010. Fast and accurate long-read alignment with Burrows–Wheeler transform. Bioinformatics 26: 589–595.

Li H, et al. 2009. The sequence alignment/map format and SAMtools. Bioinformatics 25: 2078–2079.

Lu Q, Wang K, Lei F, Yu D, Zhao H 2016. Penguins reduced olfactory receptor genes common to other waterbirds. Scientific Reports 6: 31671.

Lynch M, Force A 2000. The probability of duplicate gene preservation by subfunctionalization. Genetics 154: 459–473.

Magklara A, Lomvardas S 2013. Stochastic gene expression in mammals: lessons from olfaction. Trends in Cell Biology 23: 449–456.

Malmstrøm M, et al. 2016. Evolution of the immune system influences speciation rates in teleost fishes. Nature Genetics 48: 1204.

Malnic B, Hirono J, Sato T, Buck L 1999. Combinatorial receptor codes for odors. Cell 96: 713–723.

Matsui A, Go Y, Niimura Y 2010. Degeneration of olfactory receptor gene repertories in primates: no direct link to full trichromatic vision. Molecular Biology and Evolution 27: 1192–1200.

Miller J, et al. 2017. Hybrid assembly with long and short reads improves discovery of gene family expansions. BMC Genomics 18: 541.

Mitkus M, Nevitt G, Danielsen J, Kelber A 2016. Vision on the high seas: spatial resolution and optical sensitivity in two procellariiform seabirds with different foraging strategies. Journal of Experimental Biology 219: 3329–3338.

Mitkus M, Nevitt G, Kelber A 2018. Development of the Visual System in a Burrow-Nesting Seabird: Leach’s Storm Petrel. Brain, Behavior and Evolution 91: 4–16.

Monahan K, Lomvardas S 2015. Monoallelic expression of olfactory receptors. Annual Review of Cell and Developmental Biology 31: 721–740.

Morse DH, Buchheister CW 1977. Age and survival of breeding Leach’s storm-petrels in Maine. Bird-Banding 48: 341–349.

Nei M, Niimura Y, Nozawa M 2008. The evolution of animal chemosensory receptor gene repertoires: roles of chance and necessity. Nature Reviews Genetics 9: 951.

Nei M, Rooney A 2005. Concerted and birth-and-death evolution of multigene families. Annual Review of Genetics 39: 121–152.

Nevitt G 1999a. Foraging by seabirds on an olfactory landscape. American Scientist 87: 46–53.

Nevitt G 2000. Olfactory foraging by Antarctic procellariiform seabirds: life at high Reynolds numbers. The Biological Bulletin 198: 245–253.

Nevitt G 1999b. Olfactory foraging in Antarctic seabirds: a species-specific attraction to krill odors. Marine Ecology Progress Series 177: 235–241.

Nevitt G, Haberman K 2003. Behavioral attraction of Leach’s storm-petrels (*Oceanodroma leucorhoa*) to dimethyl sulfide. Journal of Experimental Biology 206: 1497–1501.

Nevitt G, Losekoot M, Weimerskirch H 2008. Evidence for olfactory search in wandering albatross, *Diomedea exulans*. Proceedings of the National Academy of Sciences 105: 4576–4581.

Nevitt G, Reid K, Trathan P 2004. Testing olfactory foraging strategies in an Antarctic seabird assemblage. Journal of Experimental Biology 207: 3537–3544.

Nevitt G, Veit R, Kareiva P 1995. Dimethyl sulphide as a foraging cue for Antarctic procellariiform seabirds. Nature 376: 680.

Niimura Y 2012. Olfactory receptor multigene family in vertebrates: from the viewpoint of evolutionary genomics. Current Genomics 13: 103–114.

Niimura Y 2009. On the origin and evolution of vertebrate olfactory receptor genes: comparative genome analysis among 23 chordate species. Genome Biology and Evolution 1: 34–44.

Niimura Y, Matsui A, Touhara K 2014. Extreme expansion of the olfactory receptor gene repertoire in African elephants and evolutionary dynamics of orthologous gene groups in 13 placental mammals. Genome Research 24: 1485–1496.

Niimura Y, Nei M 2005. Evolutionary dynamics of olfactory receptor genes in fishes and tetrapods. Proceedings of the National Academy of Sciences 102: 6039–6044.

O’Dwyer T, Ackerman A, Nevitt G 2008. Examining the development of individual recognition in a burrow-nesting procellariiform, the Leach’s storm-petrel. Journal of Experimental Biology 211: 337–340.

Organ C, Rasmussen M, Baldwin M, Kellis M, Edwards S. 2010. Phylogenomic approach to the evolutionary dynamics of gene duplication in birds: Wiley & Sons, New York.

Oxley J 1999. Nesting distribution and abundance of Leach’s storm-petrel (*Oceanodroma leucorhoa*) on Bon Portage Island, Nova Scotia. [Acadia University, Wolfville, Canada.

Pearce D, Hoover B, Jennings S, Nevitt G, Docherty K 2017. Morphological and genetic factors shape the microbiome of a seabird species (*Oceanodroma leucorhoa*) more than environmental and social factors. Microbiome 5: 146.

Pluznick J, et al. 2009. Functional expression of the olfactory signaling system in the kidney. Proceedings of the National Academy of Sciences 106: 2059–2064.

Pond K, Posada D, Gravenor M, Woelk C, Frost S 2006. GARD: a genetic algorithm for recombination detection. Bioinformatics 22: 3096–3098.

Pond S, Muse S. 2005. HyPhy: hypothesis testing using phylogenies: Springer, New York, NY.

Quignon P, et al. 2005. The dog and rat olfactory receptor repertoires. Genome Biology 6: R83.

Quinlan A, Hall I 2010. BEDTools: a flexible suite of utilities for comparing genomic features. Bioinformatics 26: 841–842.

Restrepo D, Lin W, Salcedo E, Yamazaki K, Beauchamp G 2006. Odortypes and MHC peptides: Complementary chemosignals of MHC haplotype? Trends in Neurosciences 29: 604–609.

Rodríguez-Ruano S, et al. 2015. The hoopoe’s uropygial gland hosts a bacterial community influenced by the living conditions of the bird. PloS One 10: e0139734.

Roper T 1999. Olfaction in birds. Advances in the Study of Behavior 28: 247.

Saito H, Chi Q, Zhuang H, Matsunami H, Mainland J 2009. Odor coding by a mammalian receptor repertoire. Science signaling 2: ra9.

Santoro S, Dulac C 2012. The activity-dependent histone variant H2BE modulates the life span of olfactory neurons. Elife 1: e00070.

Seutin G, White BN, Boag PT 1991. Preservation of avian blood and tissue samples for DNA analyses. Canadian Journal of Zoology 69: 82–90.

Simão FA, Waterhouse RM, Ioannidis P, Kriventseva EV, Zdobnov EM 2015. BUSCO: assessing genome assembly and annotation completeness with single-copy orthologs. Bioinformatics 31: 3210–3212.

Sin YW, Buesching CD, Burke T, Macdonald DW 2012. Molecular characterization of the microbial communities in the subcaudal gland secretion of the European badger (*Meles meles*). FEMS Microbiology Ecology 81: 648–659. doi: doi: 10.1111/j.1574-6941.2012.01396.x

Smit A, Hubley R, Green P 2015. RepeatMasker Open-4.0. 2013–2015. Available at www.repeatmasker.org.

Spehr M, et al. 2003. Identification of a testicular odorant receptor mediating human sperm chemotaxis. Science 299: 2054–2058.

Stamatakis A 2014. RAxML version 8: a tool for phylogenetic analysis and post-analysis of large phylogenies. Bioinformatics 30: 1312–1313.

Steiger S, Fidler A, Kempenaers B 2009a. Evidence for increased olfactory receptor gene repertoire size in two nocturnal bird species with well-developed olfactory ability. BMC Evolutionary Biology 9: 117.

Steiger S, Fidler A, Valcu M, Kempenaers B 2008. Avian olfactory receptor gene repertoires: evidence for a well-developed sense of smell in birds? Proceedings of the Royal Society B: Biological Sciences 275: 2309–2317.

Steiger S, Kuryshev V, Stensmyr M, Kempenaers B, Mueller J 2009b. A comparison of reptilian and avian olfactory receptor gene repertoires: species-specific expansion of group γ genes in birds. BMC Genomics 10: 446.

Sudmant P, et al. 2010. Diversity of human copy number variation and multicopy genes. Science 330: 641–646.

Tamura K, et al. 2011. MEGA5: molecular evolutionary genetics analysis using maximum likelihood, evolutionary distance, and maximum parsimony methods. Molecular Biology and Evolution 28: 2731–2739.

Theis K, et al. 2013. Symbiotic bacteria appear to mediate hyena social odors. Proceedings of the National Academy of Sciences 110: 19832–19837.

van-Buskirk R, Nevitt G 2008. The influence of developmental environment on the evolution of olfactory foraging behaviour in procellariiform seabirds. Journal of Evolutionary Biology 21: 67–76.

van-der-Linden C, Jakob S, Gupta P, Dulac C, Santoro S 2018. Sex separation induces differences in the olfactory sensory receptor repertoires of male and female mice. Nature Communications 9: 5081.

Vandewege M, et al. 2016. Contrasting patterns of evolutionary diversification in the olfactory repertoires of reptile and bird genomes. Genome Biology and Evolution 8: 470–480.

Warham J. 1990. The Petrels. Their Ecology and Breeding Systems. London: Academic Press.

Whittaker D, et al. 2016. Social environment has a primary influence on the microbial and odor profiles of a chemically signaling songbird. Frontiers in Ecology and Evolution 4: 90.

Wyatt TD. 2003. Pheromones and animal behaviour: communication by smell and taste. Cambridge: Cambridge University Press.

Zelano B, Edwards S 2002. An MHC component to kin recognition and mate choice in birds: predictions, progress, and prospects. The American Naturalist 160: S225–S237.

Zomer S, et al. 2009. Consensus multivariate methods in gas chromatography mass spectrometry and denaturing gradient gel electrophoresis: MHC-congenic and other strains of mice can be classified according to the profiles of volatiles and microflora in their scent-marks. Analyst 134: 114–123.

