## Supplementary material for "Olfactory receptor subgenome and expression in a highly olfactory procellariiform seabird": Table S

**Title**

**Author affiliation**

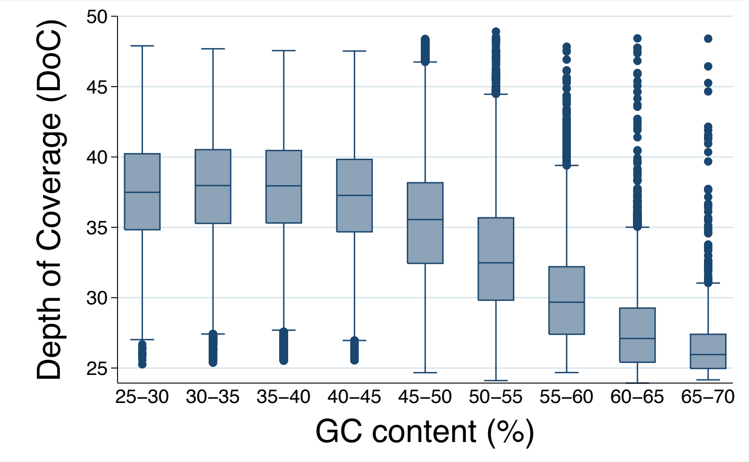


**Figure S1** The Depth of Coverage (DoC) for the bins of similar GC content.

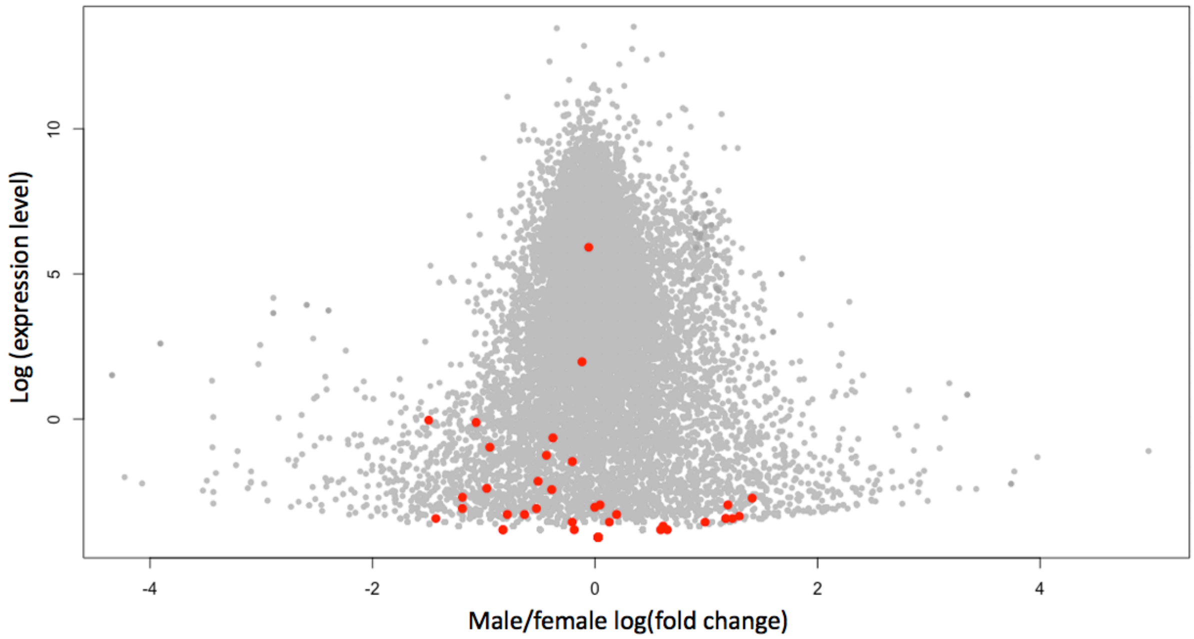


**Figure S2** Differential expression of the genes in adult male versus female olfactory epithelium. Differentially expressed genes are in dark grey. OR genes are highlighted in red. No OR genes were differentially expressed between adult males and females.

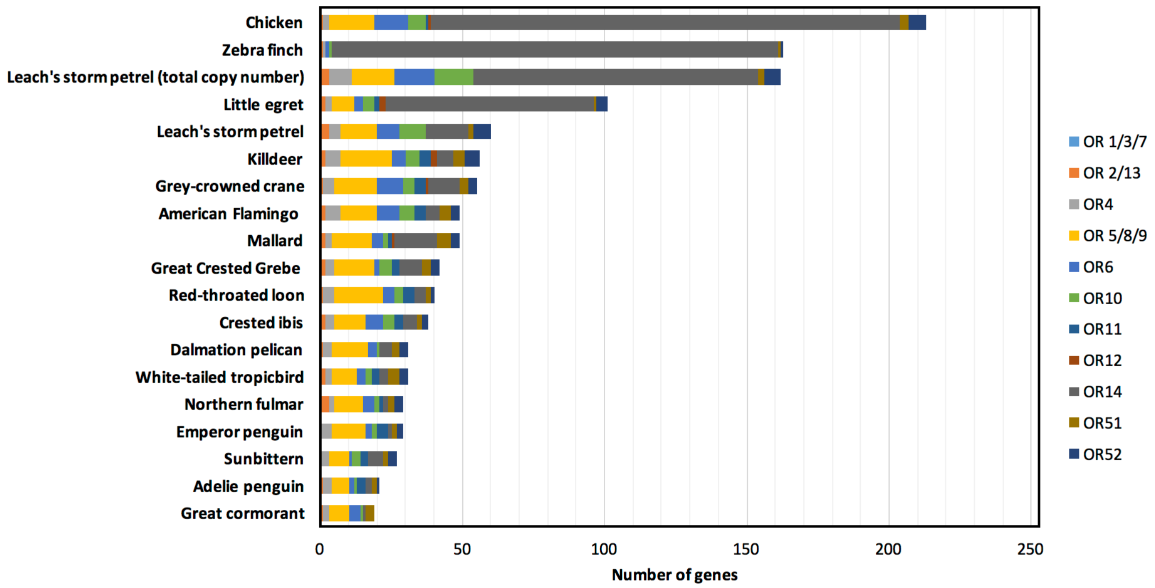


**Figure S3** The number of intact OR genes partitioned by OR gene families in waterbirds, chicken, and zebra finch. Both the OR gene number estimations based on genome annotation and copy number calculation in the Leach's storm petrel are shown. The numbers for all species except the Leach's storm-petrel are from Khan *et al*. (2015).

**Table S1** Information on tissues of Leach's storm petrel sampled in this study.

| MCZ Field No. | Sampling date | Age class | Sex | Tissue type | Dataset[A] | NCBI accession (will be provided later) |
|---|---|---|---|---|---|---|
| 15-039 | 2 Sep 2015 | Adult | F | Brain, olfactory concha | DE | |
| 15-040 | 2 Sep 2015 | Adult | M | Brain, olfactory concha, muscle, liver, heart, small intestine, kidney, tongue, eye, testes | DE, transciptome assembly, TopHat | |
| 15-041 | 2 Sep 2015 | Adult | M | Brain, olfactory concha | DE | |
| | | | | Stomach | TopHat | |
| 15-042 | 3 Sep 2015 | Chick | M? [B] | Brain, olfactory concha | DE | |
| 15-043 | 3 Sep 2015 | Adult | F | Brain, olfactory concha | DE | |
| 15-044 | 3 Sep 2015 | Adult | M | Brain, olfactory concha | DE | |
| 15-045 | 4 Sep 2015 | Chick | M? [B] | Brain, olfactory concha | DE | |
| 15-046 | 4 Sep 2015 | Adult | F | Brain, olfactory concha | DE | |
| | | | | Ovary | TopHat | |
| 15-047 | 5 Sep 2015 | Chick | M | Brain, olfactory concha | DE | |
| | | | | Spleen | TopHat | |

[A] DE: Differential expression analysis.
[B] Gonad too small to be certain.

**Table S2** Intact OR amino acid query sequences.

| Species | Species name | No. of intact OR genes | Reference |
| --- | --- | --- | --- |
| Chicken | *Gallus gallus* | 266 | Vanderwege et al. 2016 |
| Zebra finch | *Taeniopygia guttata* | 290 | Vanderwege et al. 2016 |
| American alligator | *Alligator mississippiensis* | 465 | Vanderwege et al. 2016 |
| Saltwater crocodile | *Crocodylus porosus* | 592 | Vanderwege et al. 2016 |
| Indian gharial | *Gavialis gangeticus* | 597 | Vanderwege et al. 2016 |
| Burmese python | *Python bivittatus* | 481 | Vanderwege et al. 2016 |
| Green anole | *Anolis carolinensis* | 108 | Vanderwege et al. 2016 |
| Painted turtle | *Chrysemys picta* | 842 | Vanderwege et al. 2016 |
| Chinese softshell turtle | *Pelodiscus sinensis* | 1180 | Vanderwege et al. 2016 |
| Xenopus | *Xenopus tropicalis* | 824 | Niimura 2009 |
| Zebrafish | *Danio rerio* | 154 | Niimura 2009 |
| Human | *Homo sapiens* | 388 | HORDE database |

References:

HORDE database (build #44 2 Dec 2015, accessed Oct 24, 2018)

Niimura, Y., 2009. On the origin and evolution of vertebrate olfactory receptor genes: comparative genome analysis among 23 chordate species. Genome Biology and Evolution, 1, pp.34-44.

Vandewege, M.W., Mangum, S.F., Gabaldón, T., Castoe, T.A., Ray, D.A. and Hoffmann, F.G., 2016. Contrasting patterns of evolutionary diversification in the olfactory repertoires of reptile and bird genomes. Genome biology and evolution, 8(3), pp.470-480.

**Table S3** Species included for phylogenetic analysis.

| Species | Species name | Reference |
|---|---|---|
| American alligator | *Alligator mississippiensis* | Vanderwege et al. 2016 |
| Green anole | *Anolis carolinensis* | Vanderwege et al. 2016 |
| Chicken | *Gallus gallus* | Vanderwege et al. 2016 |
| Zebra finch | *Taeniopygia guttata* | Vanderwege et al. 2016 |
| Northern fulmar | *Fulmarus glacialis* | Jarvis et al. 2014 |
| Emperor penguin | *Aptenodytes forsteri* | Jarvis et al. 2014 |
| Adelie penguin | *Pygoscelis adeliae* | Jarvis et al. 2014 |
| Great cormorant | *Phalacrocorax carbo* | Jarvis et al. 2014 |
| Crested ibis | *Nipponia nippon* | Jarvis et al. 2014 |
| Little egret | *Egretta garzetta* | Jarvis et al. 2014 |
| Dalmation pelican | *Pelecanus crispus* | Jarvis et al. 2014 |
| Red-throated loon | *Gavia stellata* | Jarvis et al. 2014 |
| White-tailed tropicbird | *Phaethon lepturus* | Jarvis et al. 2014 |
| Sunbittern | *Eurypyga helias* | Jarvis et al. 2014 |
| Killdeer | *Charadrius vociferus* | Jarvis et al. 2014 |
| Grey-crowned crane | *Balearica regulorum* | Jarvis et al. 2014 |
| Mallard | *Anas platyrhynchos* | Jarvis et al. 2014 |
| Leach's storm petrel | *Oceanodroma leucorhoa* | This study |

References:

Jarvis, E.D., Mirarab, S., Aberer, A.J., Li, B., Houde, P., Li, C., Ho, S.Y., Faircloth, B.C., Nabholz, B., Howard, J.T. and Suh, A., 2014. Whole-genome analyses resolve early branches in the tree of life of modern birds. Science, 346(6215), pp.1320-1331.

Vandewege, M.W., Mangum, S.F., Gabaldón, T., Castoe, T.A., Ray, D.A. and Hoffmann, F.G., 2016. Contrasting patterns of evolutionary diversification in the olfactory repertoires of reptile and bird genomes. Genome biology and evolution, 8(3), pp.470-480.

**Table S4** Copy number ratio of intact OR genes in the Leach's storm petrel.

| OR ID | OR family | Scaffold number | Scaffold length | Start position | End position | Copy number ratio |
|---|---|---|---|---|---|---|
| OR2-1 | OR2 | 373 | 42447 | 22343 | 23281 | 1 |
| OR2-2 | OR2 | 75 | 5450577 | 124020 | 124970 | 1 |
| OR4-1 | OR4 | 267 | 245379 | 38837 | 39763 | 1 |
| OR4-2 | OR4 | 267 | 245379 | 44800 | 45729 | 1 |
| OR4-3 | OR4 | 267 | 245379 | 49579 | 50532 | 2 |
| OR4-4 | OR4 | 300 | 133585 | 91777 | 92736 | 4 |
| OR5-1 | OR5 | 158 | 1660507 | 1639261 | 1640274 | 1 |
| OR5-10 | OR5 | 338 | 95658 | 51028 | 51960 | 2 |
| OR5-11 | OR5 | 378 | 41666 | 3678 | 4607 | 2 |
| OR5-2 | OR5 | 158 | 1660507 | 1644012 | 1644944 | 1 |
| OR5-3 | OR5 | 158 | 1660507 | 1655196 | 1656155 | 1 |
| OR5-4 | OR5 | 182 | 1123554 | 991888 | 992835 | 1 |
| OR5-5 | OR5 | 235 | 449237 | 423258 | 424226 | 1 |
| OR5-6 | OR5 | 235 | 449237 | 432355 | 433353 | 1 |
| OR5-7 | OR5 | 235 | 449237 | 438649 | 439587 | 1 |
| OR5-8 | OR5 | 267 | 245379 | 23268 | 24263 | 1 |
| OR5-9 | OR5 | 27 | 11256205 | 11240716 | 11241687 | 1 |
| OR6-1 | OR6 | 293 | 139987 | 31957 | 32907 | 1 |
| OR6-2 | OR6 | 300 | 133585 | 5445 | 6398 | 2 |
| OR6-3 | OR6 | 359 | 52130 | 38736 | 39689 | 1 |
| OR6-4 | OR6 | 409 | 29924 | 8476 | 9435 | 1 |
| OR6-5 | OR6 | 409 | 29924 | 24170 | 25111 | 1 |
| OR6-6 | OR6 | 480 | 21778 | 14483 | 15427 | 2 |
| OR6-7 | OR6 | 485 | 19036 | 201 | 1154 | 2 |
| OR6-8 | OR6 | 578 | 11286 | 3830 | 4759 | 4 |
| OR8-1 | OR8 | 158 | 1660507 | 1631284 | 1632207 | 1 |
| OR8-2 | OR8 | 235 | 449237 | 427586 | 428536 | 1 |
| OR10-1 | OR10 | 274 | 226073 | 128330 | 129277 | 4 |
| OR10-2 | OR10 | 297 | 126640 | 65519 | 66478 | 1 |
| OR10-3 | OR10 | 338 | 95658 | 76892 | 77794 | 1 |
| OR10-4 | OR10 | 345 | 61266 | 2418 | 3353 | 4 |
| OR10-5 | OR10 | 396 | 40047 | 4694 | 5626 | 1 |
| OR10-6 | OR10 | 463 | 21626 | 16656 | 17594 | 2 |
| OR10-7 | OR10 | 49 | 7629941 | 6884893 | 6885885 | 1 |
| OR10-8 | OR10 | 75 | 5450577 | 115075 | 116013 | 0 |
| OR10-9 | OR10 | 75 | 5450577 | 130763 | 131701 | 0 |
| OR13-1 | OR13 | 380 | 41223 | 13999 | 14934 | 1 |
| OR14-1 | OR14 | 263 | 246623 | 186713 | 187648 | 2 |
| OR14-10 | OR14 | 444 | 24419 | 12615 | 13547 | 1 |
| OR14-11 | OR14 | 482 | 19059 | 9009 | 9944 | 6 |
| OR14-12 | OR14 | 507 | 17189 | 13641 | 14576 | 5 |
| OR14-13 | OR14 | 513 | 16678 | 3768 | 4703 | 2 |
| OR14-14 | OR14 | 554 | 13022 | 9482 | 10417 | 45 |
| OR14-15 | OR14 | 581 | 11244 | 4964 | 5899 | 7 |
| OR14-2 | OR14 | 274 | 226073 | 53682 | 54641 | 5 |

| | | | | | | |
|---|---|---|---|---|---|---|
| OR14-3 | OR14 | 274 | 226073 | 100514 | 101449 | 3 |
| OR14-4 | OR14 | 274 | 226073 | 149030 | 149989 | 1 |
| OR14-5 | OR14 | 274 | 226073 | 170063 | 170998 | 4 |
| OR14-6 | OR14 | 274 | 226073 | 191110 | 192045 | 3 |
| OR14-7 | OR14 | 325 | 87698 | 16076 | 17011 | 3 |
| OR14-8 | OR14 | 369 | 46204 | 20875 | 21843 | 4 |
| OR14-9 | OR14 | 384 | 39901 | 16443 | 17402 | 9 |
| OR51-1 | OR51 | 205 | 852617 | 362423 | 363397 | 1 |
| OR51-2 | OR51 | 205 | 852617 | 369024 | 369980 | 1 |
| OR52-1 | OR52 | 205 | 852617 | 313130 | 314077 | 1 |
| OR52-2 | OR52 | 205 | 852617 | 324416 | 325366 | 1 |
| OR52-3 | OR52 | 205 | 852617 | 376250 | 377182 | 1 |
| OR52-4 | OR52 | 205 | 852617 | 435628 | 436578 | 1 |
| OR52-5 | OR52 | 205 | 852617 | 501313 | 502269 | 1 |
| OR52-6 | OR52 | 205 | 852617 | 528440 | 529414 | 1 |
| ORU-1 | unassigned (OR1/2/10?) | 50 | 7562026 | 6143206 | 6144225 | 1 |

**Table S5** Enriched gene ontology (GO) categories among genes significantly differentially expressed (FDR < 0.01) in the olfactory epithelium between chick and adult Leach's storm petrels. Only top 20 GO terms are shown.

| GO Term | Description | P-value | FDR q-value [A] | Enrichment [B] | N | B | n | b |
|---|---|---|---|---|---|---|---|---|
| GO:0030198 | Extracellular matrix organization | 3.03E-31 | 4.51E-27 | 12.93 | 12880 | 273 | 146 | 40 |
| GO:0043062 | Extracellular structure organization | 1.67E-30 | 1.24E-26 | 11.86 | 12880 | 305 | 146 | 41 |
| GO:0001503 | Ossification | 1.25E-16 | 6.22E-13 | 10.84 | 12880 | 99 | 288 | 24 |
| GO:0030199 | Collagen fibril organization | 1.28E-16 | 4.78E-13 | 26.39 | 12880 | 40 | 183 | 15 |
| GO:0032963 | Collagen metabolic process | 1.19E-13 | 3.54E-10 | 35.4 | 12880 | 46 | 87 | 11 |
| GO:0032502 | Developmental process | 4.71E-11 | 1.17E-07 | 1.81 | 12880 | 3881 | 193 | 105 |
| GO:0009887 | Animal organ morphogenesis | 1.22E-10 | 2.59E-07 | 5.68 | 12880 | 431 | 121 | 23 |
| GO:0001958 | Endochondral ossification | 7.05E-10 | 1.31E-06 | 17.89 | 12880 | 25 | 288 | 10 |
| GO:0036075 | Replacement ossification | 7.05E-10 | 1.17E-06 | 17.89 | 12880 | 25 | 288 | 10 |
| GO:0030282 | Bone mineralization | 2.15E-09 | 3.20E-06 | 22.44 | 12880 | 42 | 123 | 9 |
| GO:0048856 | Anatomical structure development | 6.15E-09 | 8.34E-06 | 1.96 | 12880 | 2660 | 193 | 78 |
| GO:0031214 | Biomineral tissue development | 1.21E-08 | 1.50E-05 | 15.4 | 12880 | 68 | 123 | 10 |
| GO:0007155 | Cell adhesion | 1.84E-08 | 2.11E-05 | 2.92 | 12880 | 687 | 250 | 39 |
| GO:0022610 | Biological adhesion | 2.30E-08 | 2.45E-05 | 2.9 | 12880 | 693 | 250 | 39 |
| GO:0030574 | Collagen catabolic process | 9.40E-08 | 9.34E-05 | 24.63 | 12880 | 30 | 122 | 7 |
| GO:0032964 | Collagen biosynthetic process | 1.08E-07 | 1.01E-04 | 98.7 | 12880 | 6 | 87 | 4 |
| GO:0009653 | Anatomical structure morphogenesis | 1.61E-07 | 1.41E-04 | 2.49 | 12880 | 1180 | 193 | 44 |
| GO:0070208 | Protein heterotrimerization | 3.29E-07 | 2.72E-04 | 357.78 | 12880 | 12 | 9 | 3 |
| GO:0002063 | Chondrocyte development | 4.77E-07 | 3.74E-04 | 26.83 | 12880 | 20 | 144 | 6 |
| GO:0030500 | Regulation of bone mineralization | 6.42E-07 | 4.79E-04 | 14.94 | 12880 | 57 | 121 | 8 |
| GO:0001501 | Skeletal system development | 8.15E-07 | 5.78E-04 | 13.23 | 12880 | 147 | 53 | 8 |
| GO:0003433 | Chondrocyte development involved in endochondral bone morphogenesis | 8.28E-07 | 5.61E-04 | 172.5 | 12880 | 4 | 56 | 3 |
| GO:0048513 | Animal organ development | 9.99E-07 | 6.48E-04 | 2.23 | 12880 | 1031 | 274 | 49 |
| GO:0001649 | Osteoblast differentiation | 1.74E-06 | 1.08E-03 | 4.59 | 12880 | 92 | 519 | 17 |
| GO:0071230 | Cellular response to amino acid stimulus | 2.49E-06 | 1.49E-03 | 12.51 | 12880 | 58 | 142 | 8 |
| GO:0016043 | Cellular component organization | 2.65E-06 | 1.52E-03 | 1.2 | 12880 | 4168 | 1285 | 500 |
| GO:0071840 | Cellular component organization or biogenesis | 2.88E-06 | 1.59E-03 | 1.2 | 12880 | 4209 | 1285 | 504 |
| GO:0070167 | Regulation of biomineral tissue development | 3.52E-06 | 1.87E-03 | 12.17 | 12880 | 70 | 121 | 8 |
| GO:0071229 | Cellular response to acid chemical | 4.10E-06 | 2.11E-03 | 5.48 | 12880 | 166 | 184 | 13 |
| GO:0003414 | Chondrocyte morphogenesis involved in endochondral bone morphogenesis | 4.30E-06 | 2.14E-03 | 12.31 | 12880 | 17 | 431 | 7 |
| GO:0003422 | Growth plate cartilage morphogenesis | 4.30E-06 | 2.07E-03 | 12.31 | 12880 | 17 | 431 | 7 |
| GO:0003429 | Growth plate cartilage chondrocyte morphogenesis | 4.30E-06 | 2.00E-03 | 12.31 | 12880 | 17 | 431 | 7 |
| GO:0090171 | Chondrocyte morphogenesis | 4.30E-06 | 1.94E-03 | 12.31 | 12880 | 17 | 431 | 7 |
| GO:0009888 | Tissue development | 4.35E-06 | 1.91E-03 | 10.26 | 12880 | 478 | 21 | 8 |
| GO:0043588 | Skin development | 6.50E-06 | 2.77E-03 | 18.14 | 12880 | 30 | 142 | 6 |
| GO:0050793 | Regulation of developmental process | 7.45E-06 | 3.09E-03 | 1.9 | 12880 | 2033 | 193 | 58 |
| GO:2000145 | Regulation of cell motility | 7.57E-06 | 3.05E-03 | 2.47 | 12880 | 718 | 254 | 35 |
| GO:0060536 | Cartilage morphogenesis | 1.04E-05 | 4.09E-03 | 9.2 | 12880 | 26 | 431 | 8 |
| GO:0035987 | Endodermal cell differentiation | 1.41E-05 | 5.38E-03 | 6.1 | 12880 | 25 | 844 | 10 |
| GO:0050878 | Regulation of body fluid levels | 1.76E-05 | 6.55E-03 | 2.78 | 12880 | 287 | 419 | 26 |

[A] 'FDR q-value' is the correction of the above p-value for multiple testing using the Benjamini and Hochberg (1995) method. Namely, for the ith term (ranked according to p-value) the FDR q-value is (p-value * number of GO terms) / i.

[B] Enrichment (N, B, n, b) is defined as follows:
N - is the total number of genes
B - is the total number of genes associated with a specific GO term
n - is the number of genes in the top of the user's input list or in the target set when appropriate
b - is the number of genes in the intersection
Enrichment = (b/n) / (B/N)
